# Plasmin activity and sterile inflammation synergize to promote lethal embryonic liver degeneration

**DOI:** 10.1101/2025.09.16.676624

**Authors:** Meng-Ling Wu, Courtney T. Griffin

## Abstract

Embryonic livers undergo extensive vascular expansion after midgestation to support their rapid growth and evolving functions. Immature embryonic vessels are structurally supported by extracellular matrix (ECM), which is also critical for normal liver development and function. During this same period, pro-inflammatory cytokines that function to promote hematopoiesis and hepatic organogenesis must be tightly regulated to prevent sterile inflammation. However, the contributions of endothelial cells to ECM and cytokine production during embryonic liver development are still poorly understood. Here we explore how the epigenetic chromatin-remodeling enzymes CHD4 and BRG1 work antagonistically in embryonic endothelial cells to protect developing livers from lethal degeneration. Our transcriptomic analysis of endothelial *Chd4* mutant livers, which undergo degeneration after midgestation, indicated an upregulation of both the ECM protease plasmin activity and sterile inflammation prior to the onset of lethal hepatic phenotypes. Within these pathways, we found that endothelial CHD4 and BRG1 antagonistically regulated transcription of the plasmin activator uPAR and of the inflammatory adhesion molecule ICAM-1 in developing livers. Importantly, elevated plasmin activity and sterile inflammation synergistically contribute to hepatic degeneration because a combination of genetic plasminogen reduction and treatment with the anti-inflammatory drug carprofen reduced *Chd4* mutant liver phenotypes more effectively than plasminogen deficiency or carprofen alone. Our findings highlight the critical role of endothelial cells in transcriptionally modulating plasmin activity and sterile inflammation and demonstrate the detrimental synergy of these pathways during liver development.

## INTRODUCTION

The hepatic vascular system develops in coordination with liver growth during embryogenesis (1, 2). In mouse embryos, the formation of two key hepatic vascular structures, sinusoidal capillaries and portal veins, begins around embryonic day (E) 10 (3). At this stage, the developing liver vasculature is immature and vulnerable to disruptions in the extracellular matrix (ECM), a structural component critical for maintaining vascular integrity (4–7). Liver sinusoids must maintain their structure to effectively accommodate the translocation of hematopoietic progenitor cells and blood cells between the liver parenchyma and circulation, all while preserving their barrier function. Additionally, the ECM is essential for supporting hepatic development and homeostasis (8–10). Abnormalities in the ECM can disrupt liver cell differentiation, create an imbalance in the hematopoietic niche, and trigger cellular necrosis (3, 11–16).

Another factor that compromises developing liver vasculature and contributes to lethal hepatic degeneration is excessive sterile inflammation. This conclusion emerged from observations that mouse embryos lacking components of the NFκB signaling pathway, such as p65, IKKβ, and NEMO, died with liver degeneration after midgestation (17–19). Notably deletion of the inflammatory cytokine tumor necrosis factor-α (TNFα) or its receptor TNFR1 alongside these NFκB pathway components prevented embryonic liver degeneration (18–21). These findings contributed to the current understanding that TNFα is a multifunctional cytokine, which promotes cell proliferation, differentiation, and survival during liver organogenesis in addition to facilitating hepatic inflammation and cell death under postnatal challenge conditions (22–25). Specifically, the genetic studies listed above revealed that TNFα-mediated NFκB signaling aids normal liver development in the sterile embryonic environment, although TNFα signaling results in embryonic hepatic cell death when the NFκB pathway is compromised.

ATP-dependent chromatin-remodeling complexes have emerged as pivotal regulators of vascular integrity during embryonic development, operating in spatially and temporally specific manners (26–28). These complexes utilize ATP hydrolysis to modulate chromatin accessibility, thereby shaping gene expression programs. Our previous work demonstrated that endothelial-specific deletion of CHD4 (chromodomain helicase DNA binding protein 4), the ATPase subunit within the NuRD (<u>Nu</u>cleosome <u>R</u>emodeling and <u>D</u>eacetylase) complex, leads to excessive activation of the protease plasmin, hepatic ECM degradation, sinusoidal rupture, and liver degeneration, culminating in embryonic lethality by E16.5 (27). Plasmin, which is generated from the circulating zymogen plasminogen via tissue- or urokinase-type plasminogen activators (tPA or uPA), is a key mediator of ECM remodeling and clot resolution (29). The uPA receptor (uPAR) facilitates and localizes the plasmin activation process. We found *Chd4* endothelial mutants have increased transcription of the uPAR gene (*Plaur*), which correlates with elevated hepatic plasmin activity (27). This heightened plasmin activity is consistent with our discovery of diminished expression of the ECM protein laminin, a direct target of plasmin proteolytic activity and a rich component of the basement membrane surrounding sinusoidal capillaries in embryonic livers (27). We therefore concluded that endothelial CHD4 transcriptionally inhibits excessive plasmin generation to protect the integrity of embryonic liver vasculature.

Different chromatin-remodeling complexes can act on shared genomic targets in cooperative or antagonistic ways (30–34). Like CHD4, Brahma-related gene 1 (BRG1), the ATPase in the SWI/SNF (<u>SWI</u>tch/<u>S</u>ucrose <u>N</u>on-<u>F</u>ermentable) complex, critically contributes to gene transcription in developing embryonic vasculature (28, 35, 36). Notably, BRG1 co-occupies a significant number of genomic regions with CHD4, as revealed by chromatin immunoprecipitation (ChIP) sequencing (32). We previously found that simultaneous deletion of both *Brg1* and *Chd4* in ECs allows embryonic survival until birth, despite causing defects in lung development (34). However, whether transcriptional modification of plasmin activation or of other biological processes allows *Brg1/Chd4* double mutants to bypass the lethal liver phenotypes observed in *Chd4* single mutants remains unclear.

In this study, we found that the survival of *Brg1/Chd4* double mutant embryos was transcriptionally mediated by the suppression of both aberrant plasmin activation and sterile inflammation in fetal livers. We show that *Brg1* and *Chd4* act antagonistically in regulating the transcription of key components of these pathways in ECs during liver development: *Plaur* and intercellular adhesion molecule 1 (*Icam1*). While genetic deletion of *Plg* (the gene encoding plasminogen) or treatment with the anti-inflammatory drug carprofen alone was insufficient to rescue embryonic lethality in *Chd4* mutants, co-treatment with carprofen in *Plg-*deficient *Chd4* mutants successfully improved liver degeneration and embryonic survival. These findings reveal dual transcriptional roles for ECs in regulating plasmin activity and sterile inflammation in embryonic livers, thereby highlighting the detrimental and synergistic effects of these processes on hepatic development and maintenance.

## RESULTS

### Cell death and erythrocyte accumulation seen in *Chd4-ECko* mutant livers are rescued in *Brg1/Chd4-ECdko* mutants

To understand how endothelial *Brg1/Chd4* mutants escape the lethality that occurs in endothelial *Chd4* mutant embryos by E16.5 (27, 34), we generated littermate embryos with deletions of endothelial *Chd4*, *Brg1*, or both *Brg1/Chd4* by crossing *Brg1^fl/fl^;Chd4^fl/fl^*mice with *Brg1^fl/+^;Chd4^fl/+^;VE-cadherin-Cre^+^*mice (Figure 1A). At E17.5, we detected 100% lethality in *Brg1^fl/+^;Chd4^fl/fl^;VE-cadherin-Cre^+^*(*Chd4-ECko*) embryos, 18.8% lethality in *Brg1^fl/fl^;Chd4^fl/+^;VE-cadherin-Cre^+^*(*Brg1-ECko*) embryos, and 12.5% lethality in *Brg1^fl/fl^;Chd4^fl/fl^;VE-cadherin-Cre^+^*(*Brg1/Chd4-ECdko*) embryos (Table S1). At E14.5, *Brg1/Chd4-ECdko* mutants appeared grossly comparable to control (Cre-negative) and *Brg1-ECko* embryos (Figure 1B). Notably, the histological accumulation of erythrocytes we detected in E14.5 *Chd4-ECko* mutant livers, as previously reported (27), was rescued in *Brg1/Chd4-ECdko* mutants (Figure 1C). Furthermore, we detected widespread cellular apoptosis in E14.5 *Chd4-ECko* livers by TUNEL staining and cleaved caspase-3 immunostaining, which were dramatically reduced in *Brg1/Chd4-ECdko* mutants (Figure 1D). Therefore, the lethal liver phenotypes detected in *Chd4-ECko* mutants by E17.5 were rescued in *Brg1/Chd4-ECdko* mutants.

**Figure 1.**
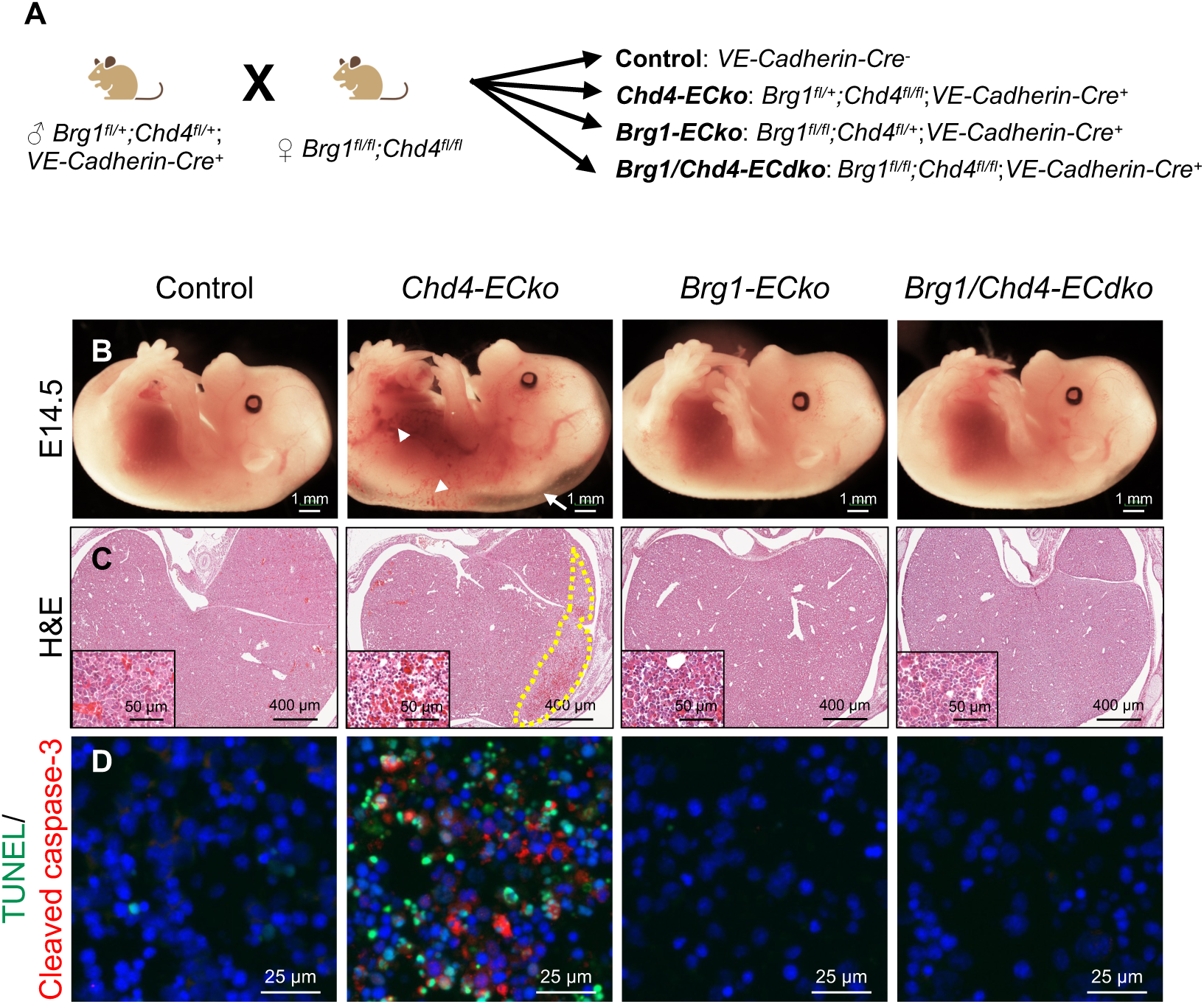
Simultaneous deletion of endothelial *Brg1* and *Chd4* rescues erythrocyte accumulation and cell death observed in *Chd4-ECko* livers. **A.** Diagram illustrating the crosses used to create endothelial-specific *Chd4*, *Brg1*, and *Brg1/Chd4* double mutant embryos. **B.** Representative images of control, *Chd4-ECko*, *Brg1-ECko*, and *Brg1/Chd4-ECdko* embryos at embryonic day (E) 14.5. N≥27 embryos for each genotype. **C.** Representative images of E14.5 control and mutant livers stained with hematoxylin and eosin (H&E). The dotted area indicates a site of liver degeneration and erythrocyte accumulation. Insets show magnified images. **D.** Representative images of E14.5 control and mutant liver sections stained with TUNEL (green) and immunostained for cleaved caspase-3 immunostaining (red). Nuclei were counterstained with Hoechst dye (blue).

Murine fetal livers become the predominant hematopoietic organ after E11.0 (3), which coincides with the time that *Chd4-ECko* embryos begin developing lethal liver phenotypes. Indeed, prior to E14.5, fetal livers are mostly comprised of erythrocytes and hematopoietic cells rather than hepatocytes (37, 38). Since some blood cells in E14.5 livers display *VE-cadherin-Cre* activity (39), we sought to rule out the possibility that the phenotypes we detected in *Chd4-ECko* livers resulted from *Chd4* deletion in the blood cell lineage. To investigate this, we crossed *Chd4-*flox mice onto the *Vav-Cre* transgenic line, which targets the hematopoietic lineage (40), and assessed embryonic lethality at E18.5. We found that *Chd4^fl/fl^; Vav-Cre^+^* embryos survived until E18.5 without evidence of mid-gestational lethality (Table S2). These results suggest that the lethal liver phenotypes seen in *Chd4-ECko* embryos are likely to be caused by *Chd4* deletion in ECs and not in hematopoietic cells.

### Plasmin deficiency does not rescue *Chd4-ECko* lethality as efficiently as endothelial *Brg1* deletion

We previously reported that deleting *Chd4* in embryonic ECs upregulates transcription of *Plaur*, thereby facilitating excessive plasmin activation, ECM degradation, and liver degeneration (27). Therefore, we sought to determine whether reduced plasmin activity contributed to the survival of *Brg1/Chd4-ECdko* embryos at E14.5. We first assessed plasmin activity by in situ zymography on fresh liver sections from E14.5 control and mutant embryos. We found the increased plasmin activity in *Chd4-ECko* livers was rescued in *Brg1/Chd4-ECdko* livers (Figures 2A and 2B). To determine if this reduction in plasmin activity correlated with reduced *Plaur* transcripts in *Brg1/Chd4-ECdko* liver ECs, we isolated LYVE1^+^ liver sinusoidal ECs (LSECs) from control and mutant livers at E12.5, two days prior to the onset of liver degeneration in *Chd4-ECko* mutants, and analyzed the samples by quantitative reverse transcription PCR (qRT-PCR). The isolated LSEC samples showed enrichment for the LSEC marker *Stab2* but not for the hepatocyte marker *Afp* (Figures S1A and S1B), and transcripts of *Chd4* and *Brg1* were reduced in LSECs from relevant mutants, as expected (Figure 2C). We found *Plaur* transcripts were upregulated in LSECs from *Chd4* mutants, downregulated in LSECs from *Brg1* mutants, and neutralized in *Brg1/Chd4-ECdko* LSECs (Figure 2C). We previously reported that *Plaur* is a direct gene target of CHD4 in a cultured murine extra-embryonic EC line (26). We questioned whether *Plaur* is also a target of BRG1 in cultured ECs. To address this question, we performed chromatin immunoprecipitation followed by quantitative PCR (ChIP-qPCR) to determine if BRG1 can interact with the *Plaur* gene promoter in the murine MS1 EC cell line. We found that BRG1 interacted with the same *Plaur* promoter region as CHD4 (−823 bp upstream of the transcription start site) (Figure 2D). Together, these findings indicate that BRG1 and CHD4 directly and antagonistically impact *Plaur* transcription in LSECs to normalize embryonic hepatic plasmin activity.

**Figure 2.**
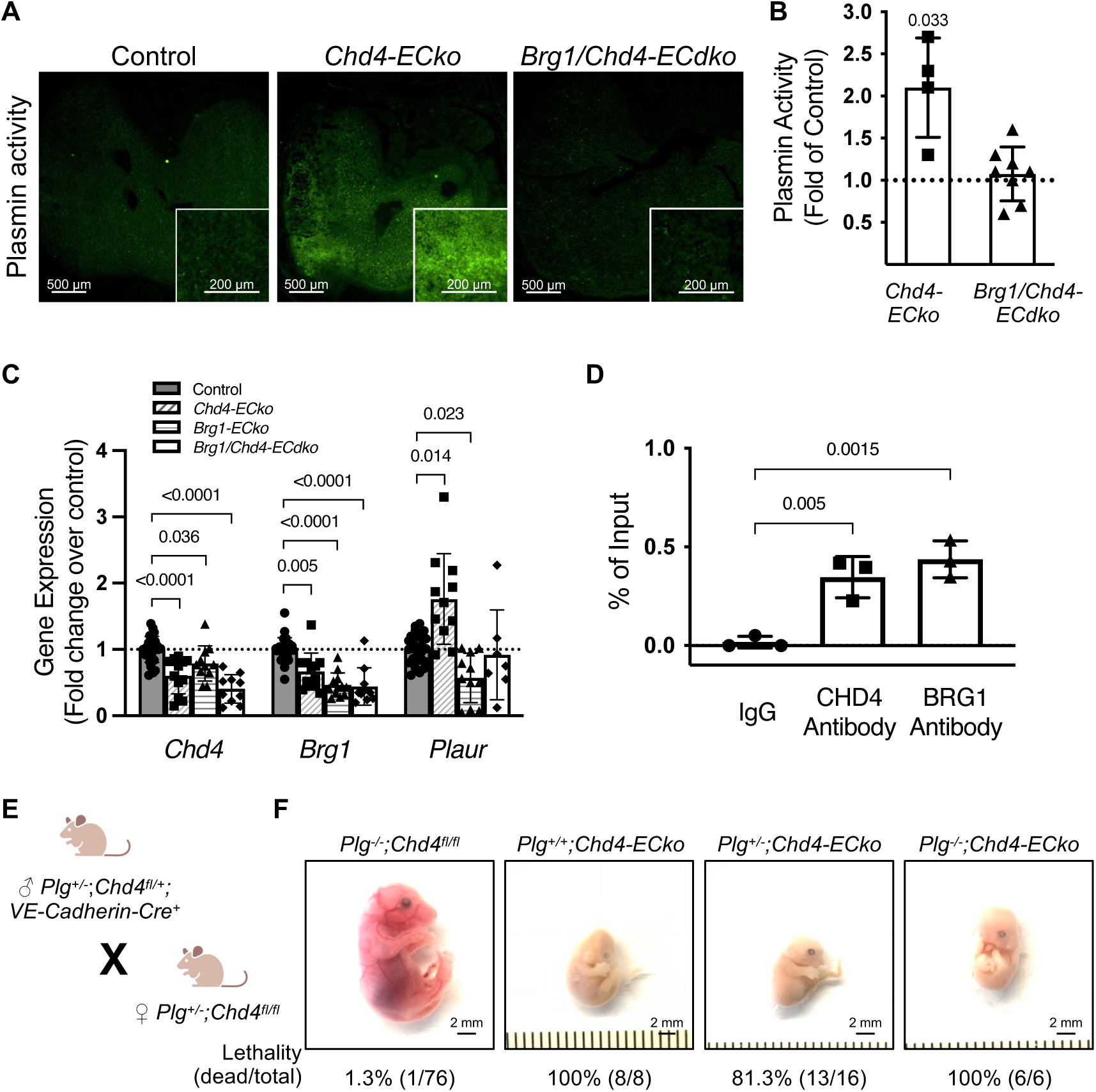
Plasmin activation observed in *Chd4-ECko* livers is rescued in *Brg1/Chd4-ECdko* embryos, but genetic plasminogen reduction does not rescue *Chd4-ECko* lethality efficiently. **A.** Representative images of plasmin activity assessed by in situ zymography on unfixed E14.5 control and mutant liver sections. Insets show magnified images. **B.** Quantification of mean fluorescence intensity of in situ plasmin activity assay as in A, normalized to relative plasmin activity of littermate controls (dotted line). N≥4 embryos. **C.** qRT-PCR analysis of *Chd4*, *Brg1*, and *Plaur* gene transcripts in liver sinusoidal ECs isolated from E12.5 control and mutant embryos. The dotted line represents the relative expression of controls. N≥7 embryos. **D.** Enrichment of CHD4 and BRG1 at the *Plaur* gene loci was assessed by ChIP-qPCR assays in the MS1 murine EC line. N=3 independent immunoprecipitations. **E.** A schematic representation of the crosses performed to generate plasminogen (*Plg*)-deficient *Chd4-ECko* embryos. **F.** Representative images and lethality rates of the indicated mutant embryos at E17.5. Data are represented as mean (±SD). 1-sample, 2-tailed t-tests were used for analysis in B. For C and D, Ordinary 1-way ANOVA with Dunnett multiple comparisons post hoc test or a Kruskal-Wallis test with Dunn multiple comparisons post hoc test was utilized when the results exhibited a nonparametric distribution.

We suspected that elevated plasmin was the primary contributor to the lethal liver phenotypes in *Chd4-ECko* livers, based on the rescue of plasmin activity, *Plaur* transcription, liver phenotypes, and lethality in *Brg1/Chd4-ECdko* mutants. Therefore, we predicted that genetic reduction of plasminogen (*Plg*), the zymogen of plasmin, in *Chd4-ECko* embryos would phenocopy the rescue seen in *Brg1/Chd4-ECdko* mutants. To test this, we crossed *Plg*^+/-^;*Chd4^fl/+^;VE-cadherin-Cre^+^* mice with *Plg*^+/-^;*Chd4^fl/fl^* mice to generate *Chd4-ECko* embryos with different doses of plasminogen deficiency (Figure 2E). To our surprise, lethality was not rescued in either *Plg*^+/-^;*Chd4-ECko* or *Plg*^-/-^;*Chd4-ECko* embryos comparably to the rescue we saw in *Brg1/Chd4-ECdko* embryos at E17.5. Thirteen out of 16 *Plg^+/-^;Chd4-ECko* embryos we dissected displayed advanced resorption at E17.5 (81.3% lethality), while each of the six *Plg^-/-^;Chd4-ECko* embryos we dissected was resorbed (100% lethality) (Figure 2F). As a reminder, this was much higher than the 12.5% lethality we saw in E17.5 in *Brg1/Chd4-ECdko* mutants (Table S1).

Coagulation abnormalities can be transferred bidirectionally between mothers and fetuses (41). Therefore, we wondered if maternal plasminogen contributed to the lethality of *Chd4-ECko* embryos with elevated uPA expression. We tested this hypothesis by generating mutant embryos from *Plg^-/-^;Chd4-ECko* dams (Figure S2A). We did see increased embryonic rescue at E17.5, with *Plg*^+/-^;*Chd4-ECko* lethality dropping to 50% and *Plg*^-/-^;*Chd4-ECko* lethality dropping to 70.8% (Figure S1B). However, this increased survival still did not match the rescue seen in *Brg1/Chd4-ECdko* embryos, indicating that other factors besides elevated plasmin likely contribute to lethal liver phenotypes in *Chd4-ECko* embryos.

### Transcriptomics reveal sterile inflammation in *Chd4-ECko* livers

To identify other potential contributors to *Chd4-ECko* lethal liver phenotypes, we performed RNA-sequencing (RNA-Seq) on CD146^+^ LSECs and non-LSEC liver cells isolated from E12.5 control and mutant livers two days before liver degeneration and lethality occurred (Figure 3A). Principal-component analysis (PCA) and clustering of EC marker genes showed distinct gene expression between LSEC and non-EC groups (Figure S3A and S3B). Because the CHD4-containing NuRD complex also contains histone deacetylases and methyl-CpG-binding domain proteins, it is typically considered a repressive complex (42). Concordantly, we found that most differentially expressed genes (DEG: log_2_ fold change > 1, adjusted p-value <0.05) in *Chd4-ECko* LSECs were upregulated compared to control LSECs (Figures 3B and S3C). Employing gene set enrichment analysis (GSEA) for control and *Chd4-ECko* LSECs, we found enrichment of “inflammatory response,” “cell activation involved in immune response,” and “TNFα signaling via NFκB” genes in *Chd4-ECko* LSECs (Figure 3C). To identify genes and pathways contributing to *Chd4-ECko* liver phenotypes, we identified 120 DEG that were exclusively changed in *Chd4-ECko* LSECs (Figure 3D). A Gene Ontology (GO) biological process analysis of these 120 DEG showed that they were associated with hallmarks of inflammation, including inflammatory and immune responses and cell activation (Figure 3E). Since TNFα signaling can mediate liver degeneration similar to what we observed in *Chd4-ECko* mutants (20), we analyzed *Tnf* transcripts from our RNA-sequencing data and found them to be significantly upregulated in *Chd4-ECko* LSECs (Figure 3F). Meanwhile, *Rela* (the gene name of NFκB p65), the effector of the NFκB pathway that counterbalances TNFα signaling to achieve normal liver development, was unaffected in all mutants (Figure S4). To analyze sterile inflammation in *Chd4-ECko* livers more directly, we next used qRT-PCR to assess transcripts of cytokine genes that were upregulated in *Chd4-ECko* LSECs by RNA-Seq. We found that *Tnf*, *Il6*, and *Il1b* transcripts were significantly increased in *Chd4-ECko* livers and were normalized in *Brg1/Chd4-ECdko* livers at E14.5 (Figure 3G). Additionally, when we exploited our RNA-Seq data to examine transcripts associated with the plasminogen activation system, we found *Plaur* was the only gene that was induced in *Chd4-ECko* LSECs at E12.5 (Figure S3D). These results indicate that in addition to promoting plasmin activation, the livers of *Chd4-ECko* embryos display evidence of heightened sterile inflammation.

**Figure 3.**
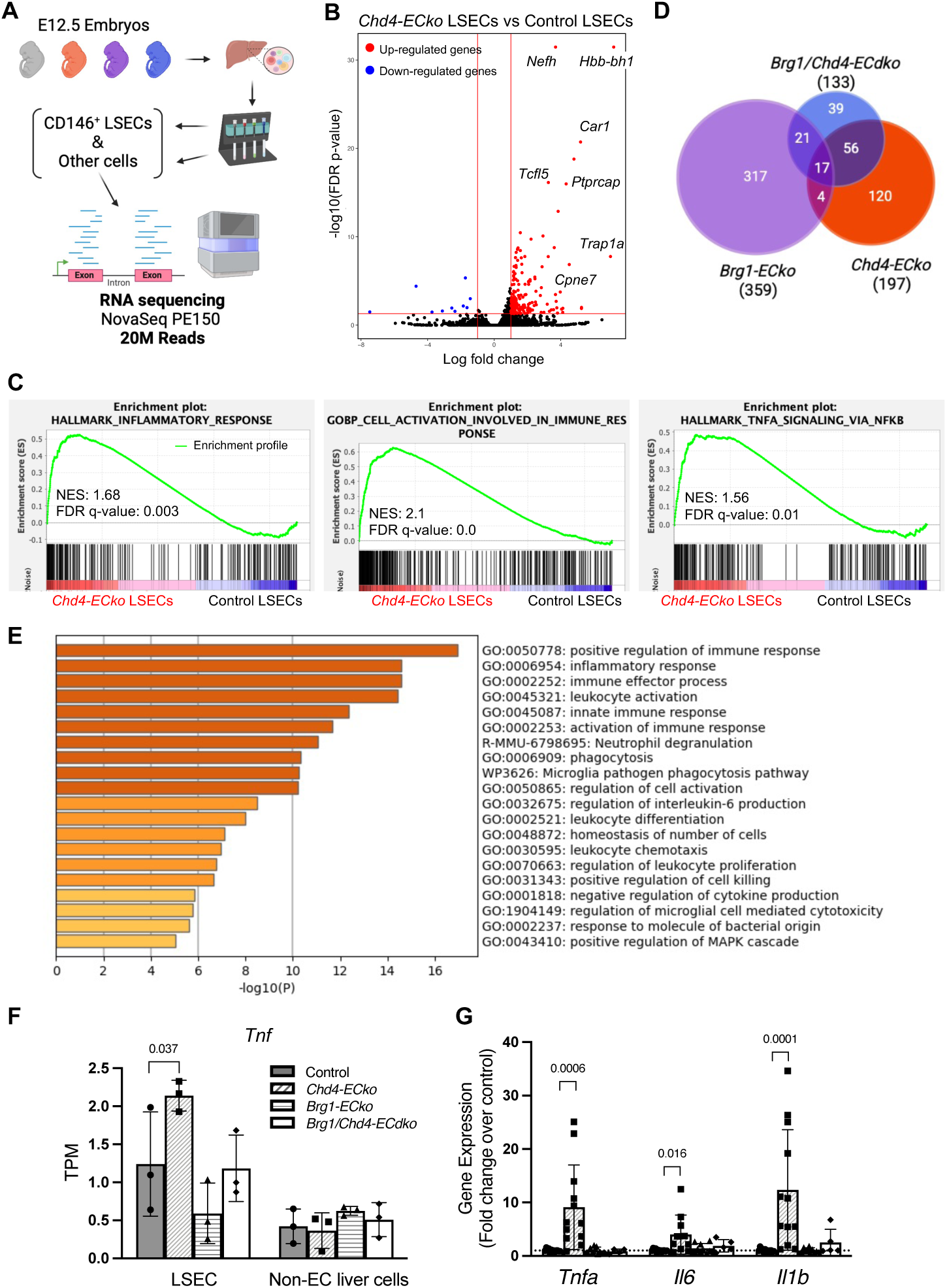
Transcriptomics indicate that *Chd4-ECko* liver sinusoidal endothelial cells (LSECs) are inflamed. **A.** A schematic diagram illustrating LSEC isolation and RNA sequencing. **B.** Volcano plot of all genes in *Chd4-ECko* LSECs compared to control LSECs. Differentially expressed genes (DEG) were defined as having more than a two-fold change in expression in *Chd4-ECko* LSECs relative to control LSECs, with an adjusted p-value <0.05. Significantly up-regulated genes: red; significantly down-regulated genes: blue. **C.** Gene set enrichment analysis (GSEA) revealed “inflammatory response,” “cell activation involved in immune response,” and “TNFα signaling via NFκB” gene sets in *Chd4-ECko* LSECs. **D.** Venn diagram illustrating the number of DEG in *Chd4-ECko*, *Brg1-ECko*, and *Brg1/Chd4-ECdko* LSECs. **E.** Gene Ontology (GO) and Kyoto Encyclopedia of Genes and Genomes (KEGG) pathway enrichment analyses were performed on the 120 DEG that were exclusively altered in *Chd4-ECko* LSECs. **F.** *Tnf* transcripts in LSECs and non-EC liver cells from E12.5 control and mutants, gleaned from RNA-sequencing data. TPM: transcripts per million mapped reads. N=3 embryos. **G.** qRT-PCR analysis of *Tnf*, *Il6*, and *Il1b* gene transcripts in E14.5 control and mutant livers. The dotted line shows the relative expression of controls. N≥4 embryos. Data are represented as mean (±SD). An ordinary one-way ANOVA with Dunnett multiple comparisons post hoc tests was used for analysis in panel F. For panel G, a Kruskal-Wallis test with Dunn multiple comparisons post hoc test was used due to the nonparametric distribution of the results.

### Expression of the endothelial adhesion molecule ICAM-1 is elevated in *Chd4-ECko* hepatic ECs

We next sought to understand endothelial factors contributing to inflammation in *Chd4-ECko* livers. Since BRG1 can bind to the *Tnf* gene locus to induce its expression in mouse kidneys after renal ischemic injury (43), we initially tried to examine BRG1 and CHD4 binding to the *Tnf* locus in control and mutant embryonic LSECs using ChIP-qPCR. However, we couldn’t collect enough LSECs from embryonic livers to perform immunoprecipitation. Instead, we investigated whether *Tnf* is a direct target of CHD4 or BRG1 in MS1 ECs using ChIP-qPCR. We found that BRG1 was enriched at the *Tnf* locus under basal conditions, but CHD4 was not enriched there (Figure S5A and S5B). Therefore, although *Tnf* was upregulated in total livers and in LSECs of *Chd4-ECko* embryos (Figure 3F and 3G), we continued our search for inflammatory factors that might be regulated directly by CHD4 in ECs.

Our RNA-Seq data revealed that transcripts for the adhesion molecule *Icam1* were upregulated in E12.5 *Chd4-ECko* LSECs and normalized in *Brg1/Chd4-ECdko* LSECs (Figure 4A). This seemed relevant to *Chd4-ECko* hepatic phenotypes because endothelial ICAM-1 induction causes leukocyte infiltration and cytokine production (44, 45) and can impair vascular barrier function (46, 47). Notably, transcripts for another endothelial adhesion molecule, *Vcam1*, were not upregulated in *Chd4-ECko* LSECs (Figure 4A). We next confirmed by qRT-PCR that *Icam1* transcripts were also elevated in *Chd4-ECko* livers at E14.5 (Figure 4B). We also assessed ICAM-1 protein expression by immunostaining and found it to be increased on both ECs and hepatoblasts of *Chd4-ECko* livers at E14.5 (Figure 4C-4E). This result was consistent with reports that cytokines can induce ICAM-1 expression in hepatocytes (48, 49). Altogether, unbiased transcriptomic data led to our discovery that *Icam1* is transcriptionally upregulated in *Chd4-ECko* embryonic livers prior to their lethal deterioration.

**Figure 4.**
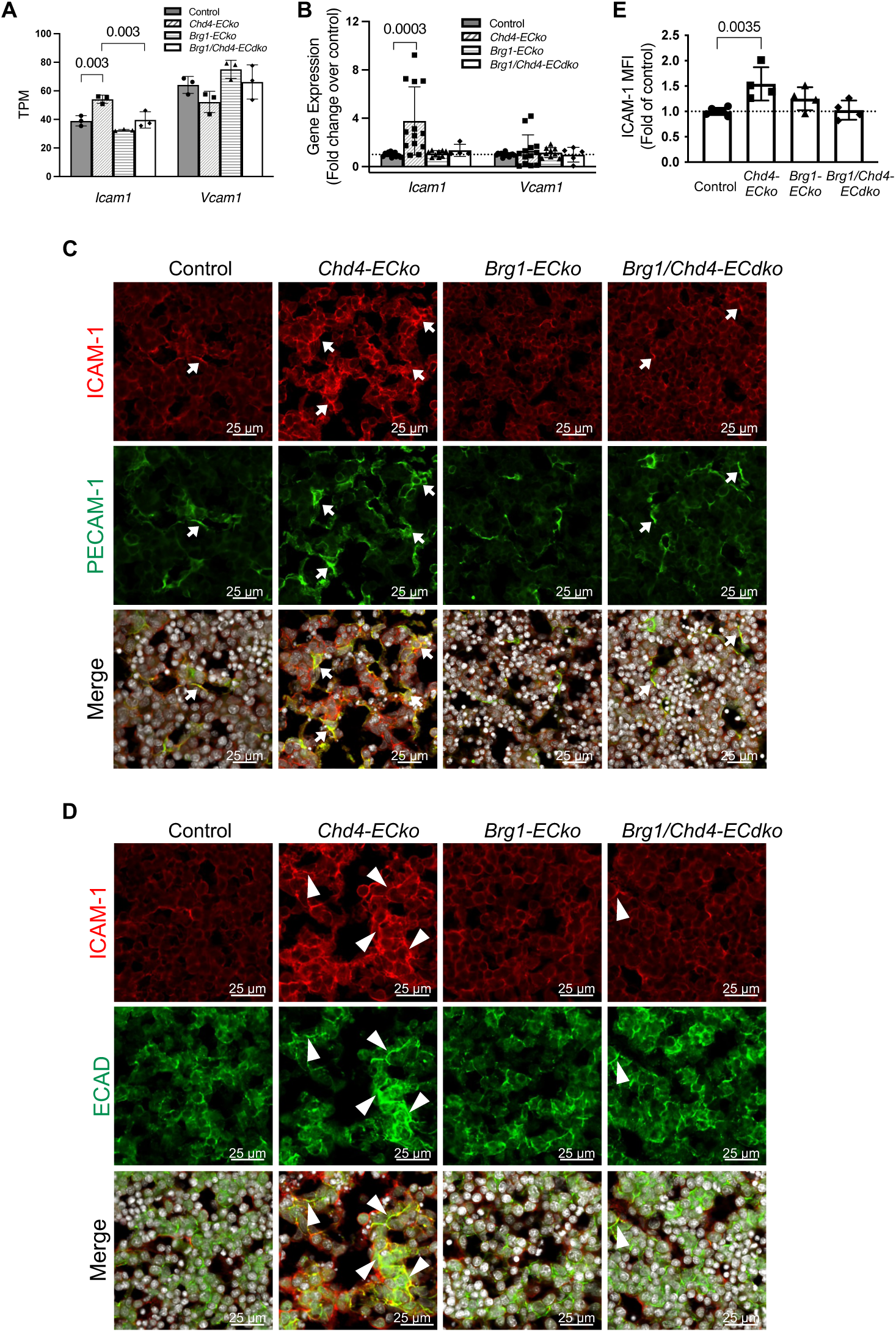
ICAM-1 expression is increased in endothelial and epithelial cells of *Chd4-ECko* livers. **A.** Transcripts of *Icam1* and *Vacm1* in E12.5 control and mutant LSECs, gleaned from RNA-sequencing data. TPM: transcripts per million mapped reads. **B.** qRT-PCR analysis of *Icam1* and *Vcam1* gene transcripts in E14.5 control and mutant livers. The dotted line shows the relative expression of controls. N≥4 embryos. **C.** Representative images of immunostaining for ICAM-1 (red) and the EC marker PECAM-1 (green) in E14.5 control and mutant livers. The merged images include a nuclear stain (Hoechst) displayed in grey. Arrows indicate cells that express both ICAM-1 and PECAM-1. **D.** Representative images of immunostaining for ICAM-1 (red) and the hepatoblast marker ECAD (green) are shown in E14.5 control and mutant livers. The merged images include a nuclear stain displayed in grey. Arrowheads indicate cells that expressed ICAM-1 and ECAD. **E.** Quantification of total ICAM-1 protein levels based on mean fluorescence intensity (MFI) in immunostained E14.5 control and mutant livers. N≥4 embryos. Data are represented as mean (±SD). The dotted line shows the relative expression of controls. Ordinary 1-way ANOVA with Dunnett multiple comparisons post hoc tests were used for analysis in A and E. For panel B, a Kruskal-Wallis test with Dunn multiple comparisons post hoc test was used due to nonparametric data distribution.

### CHD4 and BRG1 antagonistically regulate *Icam1* transcripts in ECs

BRG has been shown to bind the *ICAM1* gene promoter and promote its transcription in human umbilical vein ECs (HUVECs) (50). To investigate whether *Icam1* is also a target gene of BRG1 and CHD4 in murine ECs, we assessed *Icam1* transcripts after knocking down CHD4 and/or BRG1 in MS1 cells. We found that *Icam1* expression was upregulated upon CHD4 knockdown, downregulated upon BRG1 knockdown, and normalized when both chromatin-remodeling enzymes were knocked down simultaneously (Figure 5A). We next performed ChIP-qPCR on MS1 cells to determine whether CHD4 and BRG1 could interact directly with *Icam1* regulatory elements. We found enrichment for both CHD4 and BRG1 in the promoter region of *Icam1* (−215 bp upstream of the transcription start site) (Figure 5B). We also detected BRG1 enrichment at an intronic region (+72 bp downstream of the transcription start site) of the *Icam1* gene (Figure 5C). Collectively, these findings demonstrate that CHD4 and BRG1 work antagonistically in regulating *Icam1* expression in ECs.

**Figure 5.**
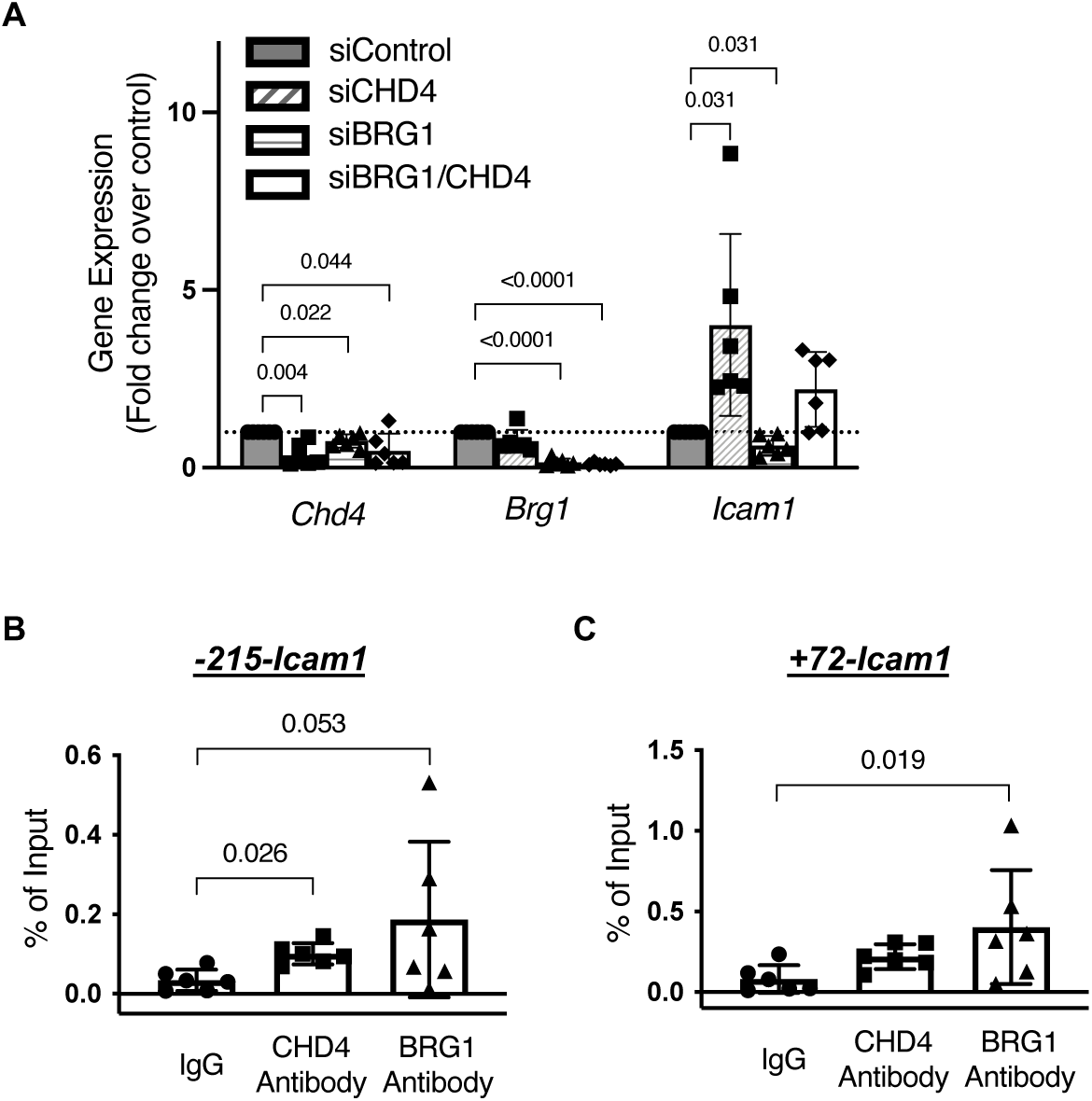
CHD4 and BRG1 antagonistically regulate *Icam1* transcription in cultured murine ECs. **A.** qRT-PCR analysis of *Chd4*, *Brg1*, and *Icam1* gene transcripts after CHD4 and/or BRG1 knockdown in MS1 ECs (n=6). The dotted line shows the relative expression in cells treated with control siRNA. **B** and **C.** ChIP-qPCR was used to determine the enrichment of CHD4 and BRG1 at regions 215 bp upstream (−215) and 72 bp downstream (+72) of the *Icam1* transcription start site in MS1 ECs. N=6 independent immunoprecipitations. Data are represented as mean (±SD). 1-sample, 2-tailed t-tests were used for analysis in A. For B and C, a Kruskal-Wallis test with Dunn multiple comparisons post hoc test was used due to nonparametric data distribution.

### Carprofen treatment combined with plasminogen deficiency rescues *Chd4-ECko* lethality

Since we observed evidence of excessive plasmin activation and inflammation in *Chd4* mutant livers, we questioned whether genetic deletion of *Plg* combined with an anti-inflammatory treatment could rescue *Chd4-ECko* liver phenotypes and lethality. We performed timed matings to generate *Chd4-ECko* embryos with *Plg* deficiency, as before, and treated pregnant dams with saline (vehicle control) or the nonsteroidal anti-inflammatory drug (NSAID) carprofen (20 mg/kg) for three consecutive days starting at E12.5 (Figure 6A). We then assessed embryonic lethality in *Plg*^+/-^;*Chd4-ECko* embryos at E17.5 since we had observed a slight improvement in *Plg*^+/-^;*Chd4-ECko* embryonic survival at baseline (81.3% lethality; Figure 2F). We found the lethality of *Plg*^+/-^;*Chd4-ECko* embryos dropped from 86% (saline-treated) to 37.5% after carprofen treatment (Figure 6B). We also assessed embryos histologically at E14.5 and found erythrocyte accumulation in saline-treated *Plg*^+/-^;*Chd4-ECko* livers was eliminated after carprofen treatment (Figure 6C and 6D). Additionally, the widespread apoptosis that we detected in saline-treated *Plg*^+/-^;*Chd4-ECko* livers was diminished after carprofen treatment (Figure 6E and 6F). Notably, treating *Chd4* mutants with carprofen alone still resulted in 89% lethality (Table 1), demonstrating that the improved survival in carprofen-treated *Plg^+/-^;Chd4-ECko* embryos was not achieved by the NSAID alone. Therefore, both plasmin activation and sterile inflammation synergistically contributed to *Chd4-ECko* embryonic liver deterioration and lethality.

**Figure 6.**
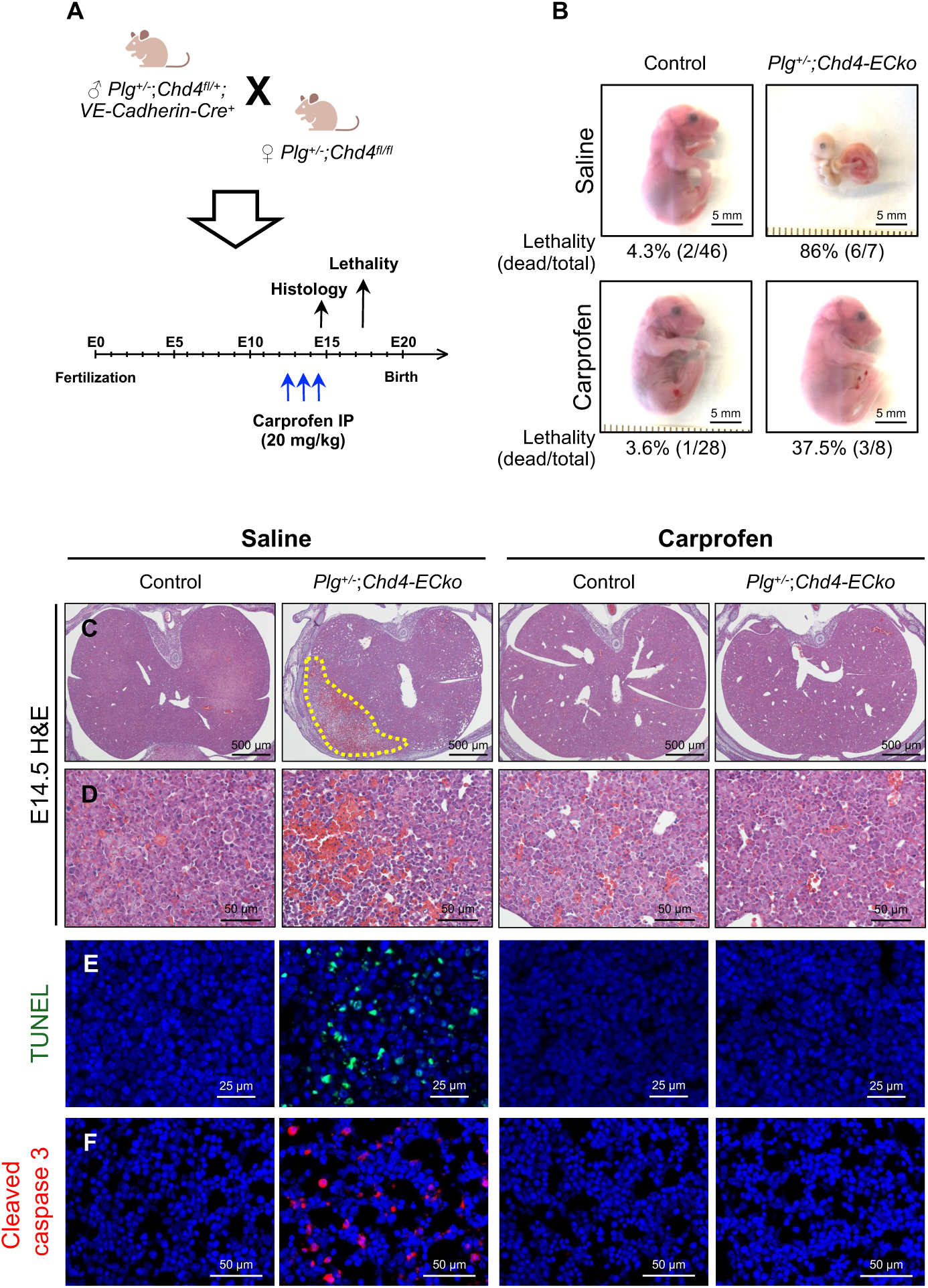
Combining anti-inflammatory carprofen treatment and genetic plasminogen deficiency rescues *Chd4-ECko* lethality efficiently. **A.** Schematic of the experimental design for the combination treatment of carprofen (20 mg/kg) and *Plg* deficiency in *Chd4-ECko* mutants. **B.** Representative images and lethality rates of control and *Plg^+/-^;Chd4-ECko* embryos after saline and carprofen treatment at E17.5. **C** and **D.** H&E staining of saline and carprofen-treated control and *Plg^+/-^;Chd4-ECko* livers at E14.5. The dotted area indicates liver deterioration and erythrocyte accumulation. **E.** Representative images of TUNEL staining (green) in saline- and carprofen-treated control and *Plg^+/-^;Chd4-ECko* liver sections at E14.5. **F.** Representative images of immunostaining for cleaved caspase-3 (red) in saline- and carprofen-treated control and *Plg^+/-^;Chd4-ECko* liver sections at E14.5. Nuclei were counterstained with Hoechst dye (blue).

**Table 1.** Lethality of carprofen-treated embryos at E17.5.

| **Genotype** | **Saline** | **Carprofen** |
| --- | --- | --- |
| Control | 0/27 (0%) | 0/35 (0%) |
| *Chd4<sup>fl/fl</sup>*; *VE-cadherin-Cre<sup>+</sup>* | 11/11 (100%) | 8/9 (89%) |

Lethality: dead/total embryos
The *P* value >0.9999 examined by Fisher's exact test

## DISCUSSION

Embryonic vessels expand to support organ growth during fetal development. Therefore, vascular defects during development can lead to cell death, organ degeneration, and embryonic lethality. This report demonstrates that endothelial chromatin remodelers CHD4 and BRG1 have antagonistic effects on the activity of the protease plasmin and on sterile inflammation in fetal livers. We found that treating with the anti-inflammatory drug carprofen, combined with plasminogen deficiency, successfully prevented liver degeneration and embryonic lethality. This indicates that ECs can play a pro-inflammatory role in embryonic livers and highlights the need for tight regulation to ensure normal liver growth.

When we began this study, we assumed that excessive plasmin activity was the primary driver of embryonic *Chd4-ECko* liver deterioration and lethality. This assumption was based on our previous report that genetic deletion of the gene encoding the plasmin activator uPA delays embryonic death in endothelial *Chd4* mutants (27). We hypothesized that loss of the plasmin precursor, plasminogen (*Plg*), would more effectively prevent death of *Chd4-ECko* mutants. However, *Plg* deficiency provided minimal rescue of *Chd4* lethality, with only 3 out of 16 *Plg^+/-^;Chd4-ECko* mutants surviving to E17.5 (81.3% lethality; Figure 2F). Interestingly, we observed a greater rescue effect when *Plg*-deficient *Chd4-ECko* mutants were generated from *Plg^-/-^* dams (Figure S2B). These findings indicate a potential role for maternal plasminogen in influencing the liver phenotypes seen in *Chd4-ECko* mutants. However, further investigation is needed to determine how maternal plasminogen crosses the placenta into fetal circulation. Notably, *Brg1/Chd4-ECko* mutants exhibited the most significant rescue of *Chd4-ECko* lethality among all the rescue effects we observed. This suggested that endothelial BRG1 and CHD4 antagonistically impacted more than just plasmin activation in embryonic livers.

While the role of the plasminogen activation system in fibrinolysis is well-recognized, there is increasing evidence that it also serves as a potent regulator of additional processes, including inflammation (29, 51–54). For example, plasmin and uPAR promote inflammation across multiple cell types by facilitating cellular activation, cytokine production, chemotaxis, cell death, and cyclooxygenase (COX)-2 production (51–60). Additionally, plasmin has been shown to induce ICAM-1 expression in HUVECs (61), and uPAR is required for leukocyte recruitment to the lung in response to *Pseudomonas aeruginosa* infection (56). In the case of uPAR, this receptor can coordinate cellular signaling through interactions with vitronectin, integrins, and growth factor receptors (29, 54, 62), so it may mediate mechanisms in addition to plasmin activation to promote inflammation. Future experiments will be needed to determine whether the elevated uPAR expression in *Chd4-ECko* mutants contributes to the sterile inflammation and liver deterioration that is not effectively rescued by *Plg* deletion alone.

Previous reports indicate that chromatin-remodeling enzymes interact in regulating inflammatory gene expression. For example, BRG1 and CHD4 antagonistically regulate the transcription of inflammatory genes in a cultured macrophage line upon lipopolysaccharide stimulation (30). Moreover, BRG1 promotes TNFα expression following liver and kidney injury (43, 63). TNFα is known to cause apoptosis and liver degeneration in mouse embryos with impaired NFκB signaling (18–20). Consistent with this evidence that unchecked TNFα signaling is detrimental to embryonic liver development, *Chd4-ECko* livers exhibited increased *Tnf* transcripts and activated TNFα signaling (Figure 3C, 3F, and 3G). However, we did not observe CHD4 interacting with the *Tnf* gene locus in the murine ECs we analyzed, so we suspect that *Tnf* is not a direct CHD4 target gene in ECs and that elevated *Tnf* expression in *Chd4-ECko* livers is an indirect consequence of *Chd4* deletion. Meanwhile, BRG1 and BRM, alternative remodeling enzymes belonging to SWI/SNF complexes, mediate the expression of ICAM-1 and VCAM-1 in ECs during the progression of atherosclerosis (50). BRG1 is reported to promote *Icam1* transcription through binding its intronic regions in both cultured ECs and glioma cell lines (50, 64). This is consistent with our observations that BRG1 is enriched in the intronic region of *Icam1* in murine MS1 ECs (Figure 5c). In this study, we provide additional evidence that CHD4 works antagonistically with BRG1 in regulating *Icam1* expression in ECs (Figure 5a). We speculate that elevated ICAM-1 expression in *Chd4-ECko* ECs may tip the hepatic environment toward detrimental sterile inflammation, thereby disrupting the normal organogenesis mediated by TNFα-triggered NFκB signaling (18–21). Nevertheless, we acknowledge that additional genetic deletion of *Icam1* or *Tnf* in *Chd4-ECko* mutants would be required to clarify the respective contributions of ICAM-1 and TNF to liver degeneration and lethality in *Chd4-ECko* embryos. Additionally, while hematopoiesis appears to be normal in p65 and IKKβ knockout livers (17, 18), ICAM-1 and inflammatory cytokines may influence hematopoiesis (65–69). Therefore, future studies could address whether hematopoiesis is dysregulated in *Chd4-ECko* mutant livers prior to lethality.

Overall, this study reveals that ECs play a crucial role in embryonic liver development through their precise transcriptional regulation of plasmin activation and sterile inflammation. These findings underscore the significant functions of ECs in ensuring proper liver development and provide insights about the synergistic and detrimental effects of plasmin and inflammation in the liver.

## MATERIALS AND METHODS

### Mouse lines and embryo dissections

*Brg1-floxed* mice (70), *Chd4-floxed* mice (71), *VE-Cadherin-Cre* transgenic mice (39), *Plg* deficient mice (72), and *Vav-iCre* transgenic mice (73) were generated and genotyped as described (31, 74–77). *Brg1^fl/fl^;Chd4^fl/fl^*mice were bred to *Brg1^fl/+^;Chd4^fl/+^;VE-Cadherin-Cre* mice to generate *Brg1-ECko, Chd4-ECko,* and *Brg1/Chd4-ECdko* mutants, which were maintained on a mixed genetic background. *Chd4^fl/fl^;Plg^+l-^*or *Chd4^fl/fl^;Plg^-l-^* mice were bred to *Chd4^fl/+^;Plg^+l-^;VE-Cadherin-Cre* mice to generate *Plg*-deficient *Chd4* mutant embryos. Mice used for timed matings ranged from eight weeks to one year of age. Noon on the day of vaginal plug detection was designated as E0.5. Embryonic yolk sacs were genotyped, and littermates that tested negative for Cre recombinase were used as controls to minimize potential genetic background effects. Saline or Carprofen (Pivetal^®^ LEVAFEN™ Injection, 20 mg/kg) was administered via intraperitoneal injection to the dams for three consecutive days starting from E12.5. Whole embryos were collected for histological analysis, and embryonic livers were isolated for indicated experiments. Mouse embryo images were captured using a Nikon SMZ800 stereomicroscope and Nikon DS-Fi1 camera. All mice were fed with a standard diet (LabDiet 5053 - PicoLab^®^ Rodent Diet 20, comprised of 20% protein and 4.5% fat) and were housed at the Oklahoma Medical Research Foundation (OMRF), which is an American Association for Accreditation of Laboratory Animal Care (AAALAC) accredited facility. Animal care was provided in accordance with the procedures outlined in the Guide for Care and Use of Laboratory Animal (National Academies Press; Washington, D.C.; 1996). All animal experiments were conducted under the approval of the OMRF Institutional Animal Care and Use Committee (IACUC). Data from male and female embryos were pooled into one group for analysis since no obvious gender effects were observed. Experimental animals were selected based on genotype; small numbers of experimental (i.e., mutant) animals within each litter excluded a need for randomization of this category.

### In situ zymography

Plasmin activity was assessed using the plasmin-specific fluorogenic substrate Boc-Glu-Lys-Lys-MCA (Peptide Institute Inc. #3105-v) as previously described (76). Freshly collected unfixed liver cryosections (12 μm) were obtained from E14.5 embryos. Sections were overlaid with in situ zymography solution containing 1% Ultra Pure LMP Agarose (Invitrogen #16520), 0.4 mM Boc-Glu-Lys-Lys-MCA, and 1 U/ml human Glu-plasminogen (Sigma-Aldrich). Overlaid sections were coverslipped and incubated for 1 hour at 37°C. To confirm that MCA cleavage was dependent on plasmin, plasminogen-free in situ zymography solution was used as a negative control. Substrate fluorescence was detected by an Eclipse Ti-E epifluorescence microscope (Nikon) and analyzed with Nikon Element (v4) or FIJI software.

### Histology and TUNEL assay

Embryos were fixed overnight at 4°C in 4% paraformaldehyde diluted in phosphate-buffered saline (PBS). After fixation, samples were dehydrated and embedded in paraffin for sectioning. Paraffin sections (4 μm) were prepared for hematoxylin and eosin (H&E) staining. Brightfield images of these sections were captured using an Eclipse 80i microscope (Nikon) and a DS-Fi1 camera (Nikon). For TUNEL staining, paraffin sections or cryosections from E14.5 control and mutant livers were analyzed with the In situ Cell Death Detection Kit, fluorescein (Roche #11684795910). In brief, paraffin sections were deparaffinized and rehydrated, while cryosections were rinsed in PBS to remove optimal cutting temperature compound (O.C.T.; Sakura Finetek). Sections were subjected to proteinase K digestion (20 μg/ml) for 30 minutes at 37°C, followed by incubation with the TUNEL reaction mixture (fluorescein-dUTP) for 1 hour at 37°C, and then rinsed three times with PBS. After TUNEL labeling, we performed immunostaining for cleaved caspase-3 and nuclear staining, if needed. Sections without TUNEL enzyme solution served as negative controls. Finally, slides were mounted using ProLong Gold mountant (Thermo Fisher Scientific #P36930). Images were acquired using an Eclipse Ti-E epifluorescence microscope or an AX-R confocal microscope (Nikon) and analyzed with Nikon Elements software (v4).

### Immunofluorescence

Embryos were dehydrated using gradients of sucrose at 10%, 15%, and 20% after fixation. They were then incubated overnight at 4°C in a 1:1 mixture of 20% sucrose and O.C.T. The following morning, embryos were embedded in O.C.T., and 8 μm sections were prepared for immunofluorescence. Sections were permeabilized with 0.2% Triton X-100 in PBS for 15 minutes and subsequently blocked with 3% BSA/PBS. All sections were immunostained overnight at 4°C with primary antibodies diluted in 1% BSA/PBS. The primary antibodies used were anti-cleaved caspase-3 (1:100, Cell Signaling #9661), anti-PECAM-1 (1:100, BD Biosciences #553370), anti-ICAM-1 (1:200, R&D SYSTEMS #AF796), and anti-ECAD (1:200, Cell Signaling #3195). Sections were washed three times with cold PBS and then incubated for 1 hour at room temperature with secondary antibodies diluted to 1:500 in 1% BSA/PBS. For nuclear counterstaining, sections were treated with 10 μg/mL Hoechst (Biotium #40046) for 5 minutes. After immunostaining, slides were mounted using ProLong Gold antifade mountant (Thermo Fisher Scientific #P36930). Images were collected using either an Eclipse Ti-E epifluorescence microscope or an AX-R confocal microscope from Nikon and were analyzed with either Nikon Elements (v4) or FIJI software.

### Liver sinusoidal endothelial cell (LSEC) isolation

LYVE1^+^ LSECs were isolated as previously described (27). E12.5 embryonic livers were digested with digestion solution (1.5 mg/mL collagenase B, 1 U/mL DNase, and 1.5 mg/mL dispase II in Dulbecco’s Modified Eagle Medium) for 15 minutes at 37°C. After digestion, cells were pelleted by centrifugation at 300 relative centrifugal force for 10 minutes at 4°C. Red blood cells were removed using 1 mL of ammonium-chloride-potassium lysing buffer (150 mmol/L ammonium chloride, 10 mmol/L sodium bicarbonate, and 1 mmol/L EDTA; pH 7.4) for 5 minutes. Cell suspension was then strained through a 70-μm cell strainer and resuspended in a mixture of Dynabeads Protein G (Invitrogen #10004D) and LYVE1 antibody (R&D SYSTEMS #AF2125) complexes, using 10 μL of beads and 1.3 μg of antibody per sample. The mixture was incubated for 30 minutes at 4°C. Immunoprecipitated LSECs and non-EC liver cells were separated with a magnetic stand and lysed in TRIzol (Invitrogen #15596026) for gene expression analysis.

### Cell culture and siRNA transfection

The MS1 adult murine pancreatic EC line (ATCC #CRL-2279) was cultured in Dulbecco’s Modified Eagle Medium supplemented with 5% fetal bovine serum. Cells were maintained in a 37°C incubator with a humidified atmosphere of 5% CO2. For siRNA-mediated gene knockdown, MS1 cells were plated in 12-well plates one day before being transfected with 50 pmol of BRG1*-*specific (Ambion #s73999), CHD4*-*specific (Ambion #s99014), or nonspecific control siRNA oligos (Ambion #AM4635). Transfections were performed using Lipofectamine RNAiMAX Transfection Reagent (Invitrogen #13778150), following the manufacturer’s protocol. Twenty-four hours after transfection, cells were serum-starved in medium containing 0.5% fetal bovine serum. RNA was extracted for gene expression analysis 48 hours post-transfection.

### RNA extraction and quantitative reverse transcription PCR (qRT-PCR)

Total RNA was extracted from livers, LSECs, or MS1 cells using Trizol, followed by purification with the RNeasy Mini Prep Kit (Qiagen #74106). RNA samples (0.1-1 μg) were converted to cDNA using the iScript cDNA Synthesis Kit (Bio-Rad #1708891). qRT-PCR analysis was performed using SsoAdvanced Universal SYBR Green Supermix (Bio-Rad #1725274) and analyzed on a CFX96 Real-Time PCR thermocycler (Bio-Rad) according to the manufacturer’s instructions. Target gene expression was normalized against at least two reference genes: *Actb*, *Gapdh*, *Rn18s*, or *Rpl13a*. The primers used were as follows: *Brg1* forward (5’-CAGTGGCTCAAGGCTATCG-3’), *Brg1* reverse (5’-TGTCTCGCTTACGCTTACG-3’); *Chd4* forward (5’-TCCTCTGTCCACCATCATCA-3’), *Chd4* reverse (5’-ACCCAAGATGGCCATATCAA-3’); *Plaur* forward (5’-GGCTTAGATGTGCTGGGAAA-3’), *Plaur* reverse (5’-CAATGAGGCTGAGTTGAGCA-3’); *Stab2* forward (5’-ATTGCTCTGGCTGCCTACTC-3’), *Stab2* reverse (5’-GTTGGCTGGCTTCTCACATC-3’); *Afp* forward (5’-AGTTTCCAGAACCTGCCGAG-3’), *Afp* reverse (5’-ACCTTGTCGTACTGAGCAGC-3’); *Tnf* forward (5’-CAGGCGGTGCCTATGTCT-3’), *Tnf* reverse (5’-AGGGTCTGGGCCATAGAACT-3’); *Il6* forward (5’-CAAAGCCAGAGTCCTTCAGAG-3’), *Il6* reverse (5’-TGGTCCTTAGCCACTCCTTC-3’); *Il1b* forward (5’-ACCAACAAGTGATATTCTCCATG-3’), *Il1b* reverse (5’-ATCCACACTCTCCAGCTGC-3’); *Icam1* forward (5’-CAATTTCTCATGCCGCACAG-3’), *Icam1* reverse (5’-AGCTGGAAGATCGAAAGTCCG-3’); *Vcam1* forward (5’-TGAACCCAAACAGAGGCAGAGT-3’), *Vcam1* reverse (5’-GGTATCCCATCACTTGAGCAGG-3’); *Actb* forward (5’-TGTTACCAACTGGGACGACA-3’), *Actb* reverse (5’-GGGGTGTTGAAGGTCTCAAA-3’); *Gapdh* forward (5’-TCAACGGCACAGTCAAGG-3’), *Gapdh* reverse (5’-ACTCCACGACATACTCAGC-3’); *Rn18s* forward (5’-CCCGAAGCGTTTACTTTGAAA-3’), *Rn18s* reverse (5’-CGCGGTCCTATTCCATTATTC-3’); *Rpl13a* forward (5’-GGATCCCTCCACCCTATGACA-3’), *Rpl13a* reverse (5’-CTGGTACTTCCACCCGACCTC-3’).

### Chromatin immunoprecipitation quantitative PCR (ChIP-qPCR)

ChIP experiments were performed using the Active Motif ChIP-IT^®^ Express Kit, following the manufacturer’s instructions. MS1 cells were cultured in 150-mm dishes without reaching full confluency. Cells were prepared by fixing the monolayer with 1% formaldehyde in serum-free media for 10 minutes at room temperature. After washing with cold PBS, 1X glycine solution was applied to cells to inhibit cross-linking. Cells were then pelleted, resuspended in lysis buffer, and incubated on ice for 15 minutes. Nuclei were isolated through Dounce homogenization (10 strokes) and subsequently pelleted by centrifugation at 500 relative centrifugal force at 4°C for 10 minutes. Pelleted nuclei were resuspended in 350 μL of shearing buffer and sonicated with a Misonix S-4000 sonicator (12 cycles of 20-second on/off pulses with an amplitude of 45) to achieve DNA fragments averaging 500 base pairs in length. Sheared chromatin was incubated with either 6 μg of IgG isotype control, anti-CHD4 (Abcam #ab70469), or anti-BRG1 antibodies (Santa Cruz #sc-17796 and Novus Biologicals #NB100-594) and rotated end-to-end at 4°C overnight. After incubation, precipitated samples were washed, eluted, and reverse cross-linked, with subsequent protein digestion performed using proteinase K. Interactions involving CHD4 and BRG1 were analyzed through qPCR. For the *Plaur* gene, the following primers were used: 5’-ACTGAGCCGCTCTGAGTGAT-3’ and 5’-CCAGGGGAAAAACAAGTTGA-3’. For the 215 bp region upstream of the *Icam1* gene (−*215-Icam1*)), the following primers were used: 5’-ACTCACCTGCTGGTCTCTGA-3’ and 5’-GCCCCTGCGATCTAGGAATTT-3’. For the 72 bp region downstream of the *Icam1* gene *(+72-Icam1*), the following primers were used: 5’-CACGCTACCTCTGCTCCTG-3’ and 5’-TGGGTCCTGGAGTTTGGAGA-3’. For the exon 1 region of the *Tnf* gene, the following primers were used: 5’-GCAGGTTCTGTCCCTTTCAC-3’ and 5’-AGTGCCTCTTCTGCCAGTTC-3’. For the exon 4 region of the *Tnf* gene, the following primers were used: 5’-TATGGCTCAGGGTCCAACTC-3’ and 5’-GCTCCAGTGAATTCGGAAAG-3’.

### LSEC RNA sequencing and analysis

LSECs were isolated from E12.5 embryonic livers as described in the LSEC isolation section, with a few modifications. Transcription inhibitors, including 5 μg/ml actinomycin D, 10 μM triptolide, and 10 μg/ml anisomycin, were added to the digestion solution (78). Additionally, CD146 MicroBeads (Miltenyi #130-092-007) were utilized to separate LSECs from non-EC liver cells. Total RNA was extracted following previously established methods, which included an on-column DNase digestion step. RNA quality was assessed using RNA ScreenTape on a TapeStation 4200 (Agilent). Library preparation and RNA sequencing (Novaseq PE150, 20 million reads) were conducted by the Clinical Genomics Center at the Oklahoma Medical Research Foundation. Repeated guanines in raw FASTQ reads from all samples were uniformly trimmed, and quality control measures were performed. Alignment to the mouse genome (GRCm39/mm39) and differential expression analysis were executed using CLC Genomics Workbench (QIAGEN). Genes exhibiting a log_2_ fold change > 1 and an adjusted p-value <0.05 when compared to LSEC-Control were classified as differentially expressed genes (DEG). Volcano plots, PCA, and heatmaps were generated using R. GSEA was performed by computing overlaps with LSEC datasets and the Molecular Signatures Database gene sets using the GSEA software tool from the Broad Institute (79). Gene Ontology (GO) enrichment analysis was conducted using the web-based portal Metascape (80).

### Statistics

Prism 10.0 software (GraphPad) was used for all statistical assessments. For in vivo experiments, each data point represents a biological replicate from a single embryo. For in vitro experiments using cultured MS1 cells, each data point represents a technical replicate from a single lot of cells. The “n” numbers in the figure legends reflect these distinctions. Statistically analyzed data are presented as mean (±SD), and associated numbers indicate adjusted p values. Data points identified as outliers by the ROUT method in Prism were excluded. Tests for normality (D’Agostino-Pearson and Shapiro-Wilk) and equal variance (Brown-Forsythe) were used to determine the appropriate parametric or non-parametric statistical model. In general, the comparison between two groups was assessed by one-sample t-tests (Figure 2B and 5A). Comparison of multiple means was made using a one-way ANOVA or a Kruskal-Wallis test, as appropriate, with relevant post hoc tests (Dunnett’s or Dunn’s, respectively).

## ACKNOWLEDGEMENTS

We thank Xie Jun, Eric Ma, Srija Nuguri, and Rahul Rajala for technical help and Griffin laboratory members for helpful discussion. We also thank the OMRF Imaging Core Facility for paraffin block embedding and the OMRF Clinical Genomics Center for technical support of RNA sequencing. Data processing and analysis were supported by the OMRF Center for Biomedical Data Sciences. We acknowledge primer sequences from PrimerBank (Massachusetts General Hospital and Harvard Medical School).

## AUTHOR CONTRIBUTION STATEMENT

Conceived and designed the experiments: M.-L.W. and C.T.G. Performed the experiments and analyzed the data: M.-L.W. Funding acquisition: M.-L.W. and C.T.G. Manuscript preparation: M.-L.W. and C.T.G.

## ETHICS STATEMENT

OMRF is an American Association for Accreditation of Laboratory Animal Care (AAALAC) accredited facility. Animal care was provided in accordance with the procedures outlined in the Guide for Care and Use of Laboratory Animal (National Academies Press; Washington, D.C.; 1996). All animal experiments were conducted under the approval of the OMRF Institutional Animal Care and Use Committee (IACUC). Data from male and female embryos were pooled into one group for analysis since no obvious gender effects were observed. Experimental animals were selected based on genotype; small numbers of experimental (i.e., mutant) animals within each litter excluded a need for randomization of this category.

## DATA AVAILABILITY STATEMENT

RNA-seq data will be made available in GEO upon acceptance of this manuscript for publication. Additional data can be requested from the corresponding author.

