## Supplemental Data for "Plasmin activity and sterile inflammation synergize to promote lethal embryonic liver degeneration"

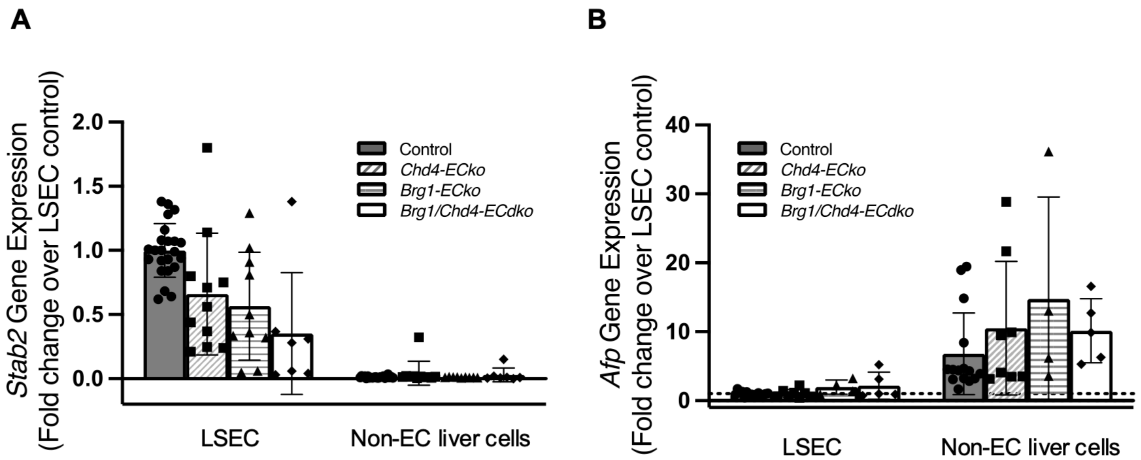


**Figure S1. Gene transcripts of liver sinusoidal endothelial cells (LSECs) and non-EC liver cells at E12.5.** LSEC marker gene *Stab2* **(A)** and hepatocyte marker gene *Afp* **(B)** levels in LSECs and non-EC liver cells from control and mutant livers. N≥4 embryos. Data are represented as mean (±SD).

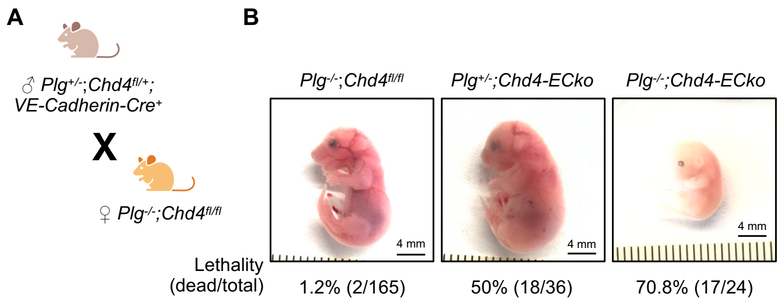


**Figure S2. Lethality of different doses of plasminogen-deficient *Chd4-ECko* embryos generated from plasminogen null dams. A.** A schematic representation of the crosses used to generate plasminogen (*Plg*)-deficient *Chd4-ECko* embryos from *Plg* null dams is shown. **B.** Representative images and the lethality rate of control, *Plg*$^{+/-}$*;Chd4-ECko*, and *Plg*$^{-/-}$*;Chd4-ECko* embryos at E17.5.

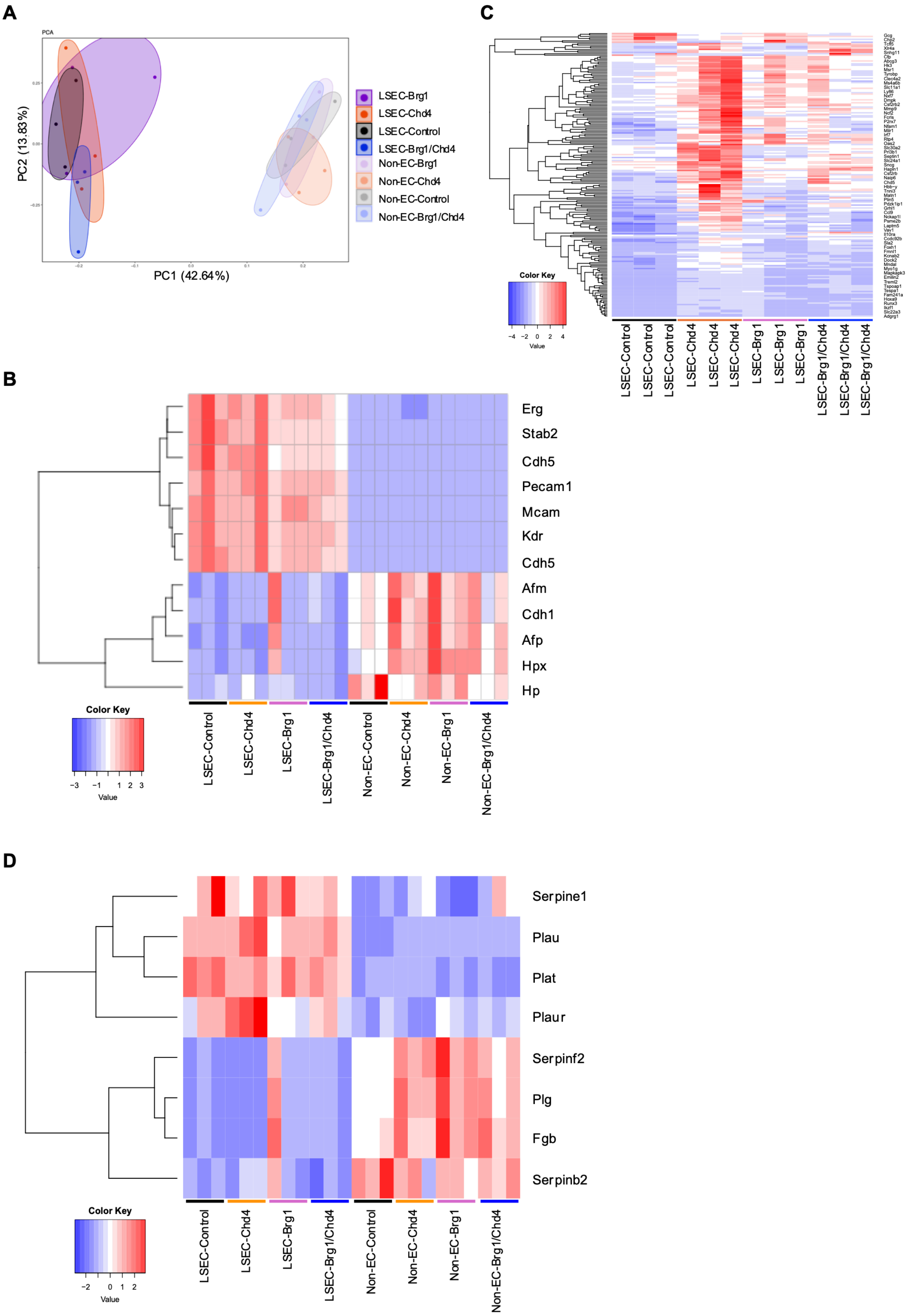
A
PCA
PC1 (42.64%)
PC2 (13.83%)
LSEC-Brg1
LSEC-Chd4
LSEC-Control
LSEC-Brg1/Chd4
Non-EC-Brg1
Non-EC-Chd4
Non-EC-Control
Non-EC-Brg1/Chd4
B
Erg
Stab2
Cdh5
Pecam1
Mcam
Kdr
Cdh5
Afm
Cdh1
Afp
Hpx
Hp
Color Key
Value
LSEC-Control
LSEC-Chd4
LSEC-Brg1
LSEC-Brg1/Chd4
Non-EC-Control
Non-EC-Chd4
Non-EC-Brg1
Non-EC-Brg1/Chd4
C
Color Key
Value
LSEC-Control
LSEC-Control
LSEC-Control
LSEC-Chd4
LSEC-Chd4
LSEC-Chd4
LSEC-Brg1
LSEC-Brg1
LSEC-Brg1
LSEC-Brg1/Chd4
LSEC-Brg1/Chd4
LSEC-Brg1/Chd4
D
Serpine1
Plau
Plat
Plaur
Serpinf2
Plg
Fgb
Serpinb2
Color Key
Value
LSEC-Control
LSEC-Chd4
LSEC-Brg1
LSEC-Brg1/Chd4
Non-EC-Control
Non-EC-Chd4
Non-EC-Brg1
Non-EC-Brg1/Chd4

**Figure S3. Transcription profiles from RNA-sequencing of LSEC and non-EC samples from control and mutant livers at E12.5. A.** Principal component analysis (PCA) of RNA-sequencing samples from LSEC and non-EC groups, with outputs from 18,241 genes. **B.** Heatmap illustrating gene expression of EC and hepatic cell markers in LSEC and non-EC samples. **C.** Heatmap showing differentially expressed genes (DEG) in LSECs between control and *Chd4-ECko* LSECs. DEG were defined as having more than a two-fold change in expression in *Chd4-ECko* LSECs relative to control LSECs, with an adjusted p-value <0.05. **D.** Heatmap showing gene expression of plasmin activation pathway molecules in LSEC and non-EC samples.

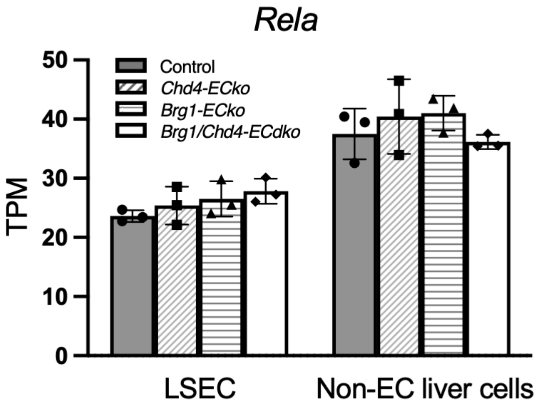


**Figure S4. *Rela* transcripts in LSECs and non-EC liver cells at E12.5.** *Rela* transcripts gleaned from RNA-sequencing data. TPM: transcripts per million mapped reads. Data are represented as mean (±SD).

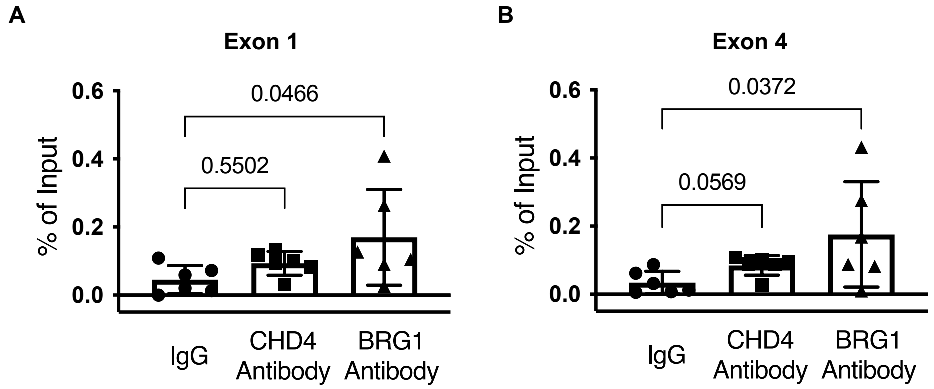


**Figure S5. BRG1 interacts with the *Tnf* gene locus in cultured MS1 cells.** ChIP-qPCR was used to determine the enrichment of CHD4 and BRG1 at exon 1 (**A**) and exon 4 (**B**) of the *Tnf* gene locus in MS1 ECs. N=6 independent immunoprecipitations. Data are represented as mean (±SD). Ordinary 1-way ANOVA with Dunnett multiple comparisons post hoc tests was used for analysis in A. For B, a Kruskal-Wallis test with Dunn multiple comparisons post hoc test was used due to nonparametric data distribution.

**Table S1**

**Lethality of endothelial mutants at E17.5**

(from *Brg1*$^{fl/fl}$*;Chd4*$^{fl/fl}$ x *Brg1*$^{fl/+}$*;Chd4*$^{fl/+}$*;VE-cadherin-Cre*$^{+}$)

| Genotype | Observed embryos (%) | Resorptions | Lethality (dead/total) |
|---|---|---|---|
| Control | 63 (75.9%) | 0 | 0% |
| *Brg1*$^{fl/+}$*;Chd4*$^{fl/fl}$*;VE-cadherin-Cre*$^{+}$ (*Chd4-ECko*) | 0 (0%) | 11 | 100% |
| *Brg1*$^{fl/fl}$*;Chd4*$^{fl/+}$*;VE-cadherin-Cre*$^{+}$ (*Brg1-ECko*) | 13 (15.7%) | 3 | 18.8% |
| *Brg1*$^{fl/fl}$*;Chd4*$^{fl/fl}$*;VE-cadherin-Cre*$^{+}$ (*Brg1/Chd4-ECdko*) | 7 (8.4%) | 1 | 12.5% |
| Total | 83 | | |

**Table S2**

**Embryonic genotypes at E18.5**

(*Chd4*$^{fl/+}$;*Vav-Cre*$^{+}$ X *Chd4*$^{fl/fl}$)

| **Genotype** | **Observed embryos** | **Expected embryos** |
|---|---|---|
| Control | 23 | 20 |
| *Chd4*$^{fl/+}$;*Vav-cre*$^{+}$ | 8 | 10 |
| *Chd4*$^{fl/fl}$;*Vav-cre*$^{+}$ | 9 | 10 |
| Total | 40 | |

$X^2$ = 0.48 with 2 degrees of freedom.
